# Comparative Genomic Insights into the Evolution of Aquatic and Terrestrial Adaptations in Plants

**DOI:** 10.64898/2026.04.02.716071

**Authors:** Sébastien Cabanac, Christophe Dunand, Catherine Mathe

## Abstract

Terrestrial plants emerged from the water about 500 million years ago. Thereafter, they have diversified and now inhabit most of the Earth’s surface. More recently, some species have re-adapted to an aquatic lifestyle, both in fresh and salt water, and fully or partially submerged. The mechanisms enabling these adaptations between terrestrial and aquatic life are extremely numerous, making it difficult to have a comprehensive overview of the phenomenon. Here, we performed a series of intraspecific measurements of the selection pressure affecting orthologous genes in eight aquatic and four terrestrial plants. Our analyses showed that aquatic plants have a relaxed selection pressure on nutrient assimilation mechanisms, probably linked to a greater bioavailability, as well as stronger adaptations to oxidative stress, while terrestrial plants evolution is linked to environment perception. Inter-species analyses have also highlighted a different evolution of chloroplast proteins between these two types of plants, suggesting adaptations to gas availability.

## Introduction

Plants have emerged from water around 450-500 million years ago [1]. Their terrestrialization required significant adaptations to cope with a radically different and more stressful environment [2]. In particular, plants had to adapt to much more intense UV radiation, gravity, temperature variations, desiccation as well as changes in the availability of nutrients, oxygen and CO_2_. While most of the stresses experienced by plants during such events are obvious, the mechanisms of adaptation are numerous and not yet fully identified. Conversely, today, many plant species distributed among the entire land plant clade have independently re-adopted an aquatic lifestyle [3,4]. It can be suggested that these plants have lost or are in the process of losing the mechanisms of adaptation to a terrestrial lifestyle, while “re-”adapting to an aquatic lifestyle, notably reduced oxygen availability [5].

Adaptations to different lifestyles may be achieved through genetic adaptations, and it can be assumed that orthologous genes that are essential for aquatic lifestyle have undergone convergent adaptation in some, or all, aquatic plants. These genes should be subject to less purifying selection in terrestrial plants, and conversely genes that are essential for the terrestrial lifestyle should be subject to stronger selection pressure in terrestrial plants than in aquatic ones. To study the difference in selection between terrestrial plant genes and their aquatic plant orthologs, we used genetic variability data for a set of eight aquatic and four terrestrial species. The aquatic species include *Lemna minor, Nelumbo nucifera, Nymphoides peltata, Pontederia crassipes, Posidonia oceanica, Ranunculus bungei, Sparganium stoloniferum* and *Zostera marina*. The terrestrial plant species include *Arabidopsis thaliana, Brachypodium distachyon, Marchantia polymorpha* and *Medicago truncatula*. We also compared intra-species variability of orthologous genes among a set of 19 aquatic and 32 terrestrial species, taking species representing all the angiosperms and associating each aquatic plant with close terrestrial species when possible. We further took advantage of the large amount of available chloroplast genomes to study this same type of evolution on an even larger number of species, including 33 aquatic species and 45 terrestrial species.

## Results

### Comparison of selection pressures between aquatic and terrestrial plants

Comparison of selection pressure between orthologs of eight aquatic plants (*L. minor, N. nucifera, N. peltata, P. crassipes, P. oceanica, R. bungei, S. stoloniferum, Z. marina*) and four terrestrial plants (*A. thaliana, B. distachyon, M. polymorpha, M. truncatula*) indicates that the orthogroups that best contribute to the distinction between aquatic and terrestrial lifestyles were generally more conserved in terrestrial plants: indeed, an average Tajima’s D [6] value of −0.33 is obtained for terrestrial plants and 0.25 for aquatic ones (Figure S1). This observation is also confirmed when Tajima’s D values are examined independently for each species, including *M. polymorpha*, indicating that the results represent a general trend among angiosperms and possibly among all embryophytes (Figure 1).

**Figure 1.**
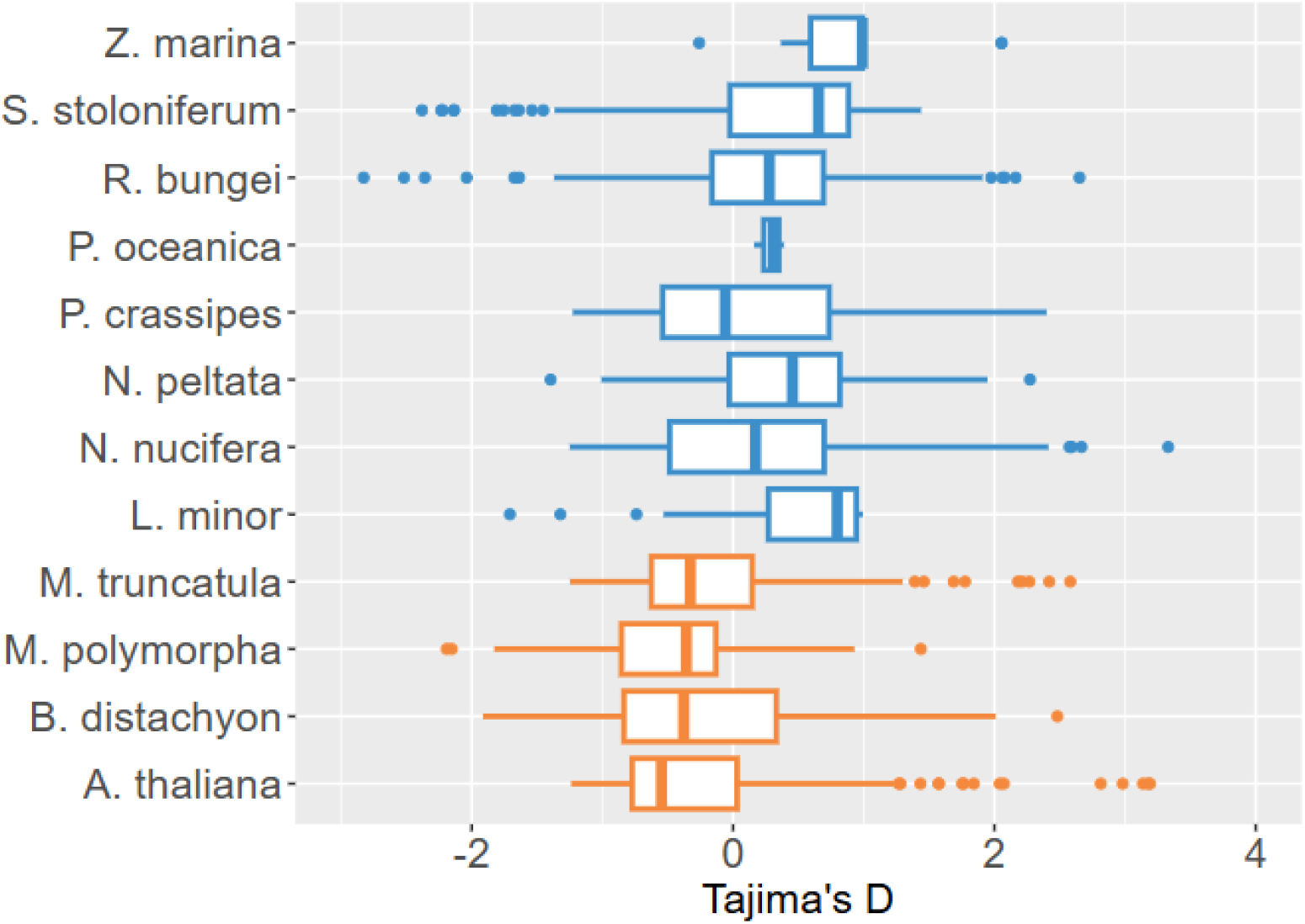
Distribution of scaled Tajima’s D among the orthologs that best explained the distinction between aquatic and terrestrial lifestyles. Lower Tajima’s D indicates stronger background selection. Orange boxplots are the results for terrestrial species and blue boxplots for aquatic species.

Selective sweep, balanced selection and population effect hypotheses are unlikely, as we also calculated a slightly negative average Zeng’s E (−0.37) [7] and an almost null Way & Fu’s H (0.03) (Figure S1) [8] on terrestrial species. This weak background selection that we found allows a good separation of the two groups of species by a sparse Projection to Latent Structure Discriminant Analysis (sPLS-DA), indicating that genetic variability can explain the differences between the two ways of life (Figure S2).

Enrichments in GO terms, KEGG pathways [9,10], Reactome pathways, COMPARTMENTS’ subcellular localizations and UniProt’s keywords were carried out on *A. thaliana* genes within each of the orthogroups that best explained the distinction between aquatic and terrestrial lifestyles (Table 1). Different enrichments were performed for orthogroups with stronger background selection in aquatic plants (Table S1) or terrestrial plants (Table S3).

**Table 1.** Enriched GO terms in *A. thaliana* obtained from genes among the orthogroups that best explained the distinction between aquatic and terrestrial lifestyles. Lifestyle column indicate which kind of plants have a strongest conservation (*i*.*e*. lowest average Tajima’s D) for each GO term. For clarity, only GO terms with REVIGO-calculated dispensability lower than 0.1 are shown and uninformative GO terms (*e*.*g*. metabolic process) were manually removed.

| Term ID | Description | FDR | Lifestyle |
| --- | --- | --- | --- |
| GO:0050113 | inositol oxygenase activity | 8.54E-07 | Aquatic |
| GO:0006020 | inositol metabolic process | 1.54E-06 | Aquatic |
| GO:0005506 | iron ion binding | 3.80E-04 | Aquatic |
| GO:2000377 | regulation of reactive oxygen species metabolic process | 7.30E-04 | Aquatic |
| GO:0006470 | protein dephosphorylation | 8.80E-04 | Terrestrial |
| GO:0047372 | monoacylglycerol lipase activity | 1.10E-03 | Terrestrial |
| GO:0010618 | aerenchyma formation | 3.00E-03 | Aquatic |
| GO:0010617 | circadian regulation of calcium ion oscillation | 4.10E-03 | Aquatic |
| GO:0012501 | programmed cell death | 4.10E-03 | Aquatic |
| GO:0015137 | citrate transmembrane transporter activity | 4.40E-03 | Aquatic |
| GO:0071585 | detoxification of cadmium ion | 7.20E-03 | Terrestrial |
| GO:2000779 | regulation of double-strand break repair | 7.20E-03 | Terrestrial |
| GO:0009882 | blue light photoreceptor activity | 1.25E-02 | Aquatic |
| GO:0042393 | histone binding | 1.80E-02 | Terrestrial |
| GO:0005488 | binding | 4.06E-02 | Terrestrial |
| GO:0019888 | protein phosphatase regulator activity | 4.94E-02 | Terrestrial |

To take into account non-synonymous substitutions, all the analyses carried out from Tajima’s D values were also done with the ratio of the number of non-synonymous polymorphic sites to the number of synonymous polymorphic sites (NS/S) for each gene (Table 2, Figure S1, Figure S2b, Figure S2c, Table S2, Table S4).

**Table 2.** Enriched GO terms in *A. thaliana* obtained from genes among the orthogroups that best explained the distinction between aquatic and terrestrial lifestyles. Lifestyle column indicates which kind of plants have a strongest conservation (*i*.*e*. lowest average NS/S) for each GO term. For clarity, only GO terms with REVIGO-calculated dispensability lower than 0.1 are shown and uninformative GO terms (*e*.*g*. metabolic process) were manually removed.

| Term ID | Description | FDR | Lifestyle |
| --- | --- | --- | --- |
| GO:0009734 | auxin-activated signaling pathway | 2.14E-17 | Terrestrial |
| GO:0030234 | enzyme regulator activity | 1.64E-13 | Aquatic |
| GO:0031399 | regulation of protein modification process | 1.19E-11 | Aquatic |
| GO:0050105 | L-gulonolactone oxidase activity | 2.79E-10 | Aquatic |
| GO:0019853 | L-ascorbic acid biosynthetic process | 3.07E-08 | Aquatic |
| GO:0052324 | plant-type cell wall cellulose biosynthetic process | 5.42E-06 | Terrestrial |
| GO:0046524 | sucrose-phosphate synthase activity | 2.51E-05 | Terrestrial |
| GO:0042910 | xenobiotic transmembrane transporter activity | 6.91E-05 | Terrestrial |
| GO:0050896 | response to stimulus | 5.70E-04 | Terrestrial |
| GO:0071949 | FAD binding | 9.30E-04 | Aquatic |
| GO:0005488 | binding | 9.30E-03 | Terrestrial |
| GO:0005249 | voltage-gated potassium channel activity | 2.05E-02 | Aquatic |
| GO:0071836 | nectar secretion | 2.34E-02 | Terrestrial |
| GO:0010244 | response to low fluence blue light stimulus by blue low-fluence system | 3.77E-02 | Aquatic |
| GO:0022414 | reproductive process | 3.77E-02 | Aquatic |
| GO:0032502 | developmental process | 4.06E-02 | Aquatic |

An enrichment of GO terms associated with inositol is found in aquatic plants, and is mainly linked to *Myo-Inositol Oxygenases (MIOXs)*. Those genes are involved in the synthesis of ascorbate [11], a major antioxidant involved in ROS scavenging [12]. The GO term “Ascorbate and aldarate metabolism” is also highly enriched (Table S2), and is linked to the *GULLO1-7* genes [13], and to two ascorbate peroxidases. An enrichment in EDS1 disease-resistance complex is also obtained (Table S1), linked in particular to the presence of *PAD4* and *EDS1*. These genes, along with others, are also involved in the enrichment of other terms, such as “Regulation of hydrogen peroxide metabolic process”, “Programmed cell death” and “Aerenchyma formation”. Terms related to pectin catabolism were also enriched, which is essential for aerenchyma formation [14]. “Cryptochrome, plant” enrichment is linked to *CRY1* and *CRY2* (Table S1). They encode photoreceptors that perceive blue light and are involved in the circadian cycle, and in a large set of biological processes regulating the development of the plant [15]. Interestingly, a set of genes potentially involved in cryptochromes-dependent growth regulation was also found (*e*.*g. EBF2, PIF5, PIF4, EBF1*, see Table S2).

In contrast, the enriched GO terms for terrestrial plants contain many terms such as “Late endosome membrane”, “Intracellular transport”, “Dynamin GTPase effector domain” (Table S3), “Antiporter activity”, “Signal transduction”, “Cell communication”, “Response to chemical”, “Cellular response to stimulus”, “Protein kinase activity” and “Response to stimulus” (Table S4), which suggest an increased and rapid perception of stimuli and interactions with the external and cellular environments. Part of these enrichments is due to the presence of numerous IAA-related genes, which also allows a strong enrichment of the GO term “Auxin-activated signaling pathway” (Table S4). Other genes associated with stronger selection pressures in terrestrial plants seems to be linked to nutrition. They are notably enriched in “Patatin-like phospholipase”, APR-related term “Sulfate assimilation”, and autophagy-under-starvation-related terms (Table S3). Patatin-like genes regulate root architecture in case of phosphate deficiency [16], while *APR1, APR2* and *APR3* have important role in the regulation of sulfate assimilation [17]. GO terms related to the cell wall are also found with terrestrial plants, but are rather associated with cellulose synthesis and are linked to the presence of COBRA-like proteins.

Some orthogroups appear particularly important in explaining the distinction between the two lifestyles, which is evidenced by their high absolute values of sPLS-DA’s “loadings”. Notably, a *DTX-*related orthogroup with stronger background selection is found in terrestrial plants. Conversely, an *AKT-*related orthogroup is more conserved in aquatic plants. This orthogroup contains AKT genes involved in potassium transport [18].

### Inter-species selection pressure

We used codeML to compute the selective pressure ratio ω of orthologs from the nuclear genomes of 19 aquatic angiosperms and 32 terrestrial angiosperms (Figure S3). We also did this for a larger set of chloroplastic genomes (Figure S4). Significantly different ω between aquatic and terrestrial plants were found in only three orthogroups from the nuclear genome, and seven from the chloroplast genome. The functions of the genes from the nuclear genome were generally poorly understood, contrasting with the seven orthogroups identified from chloroplastic genomes (Figure 2).

**Figure 2.**
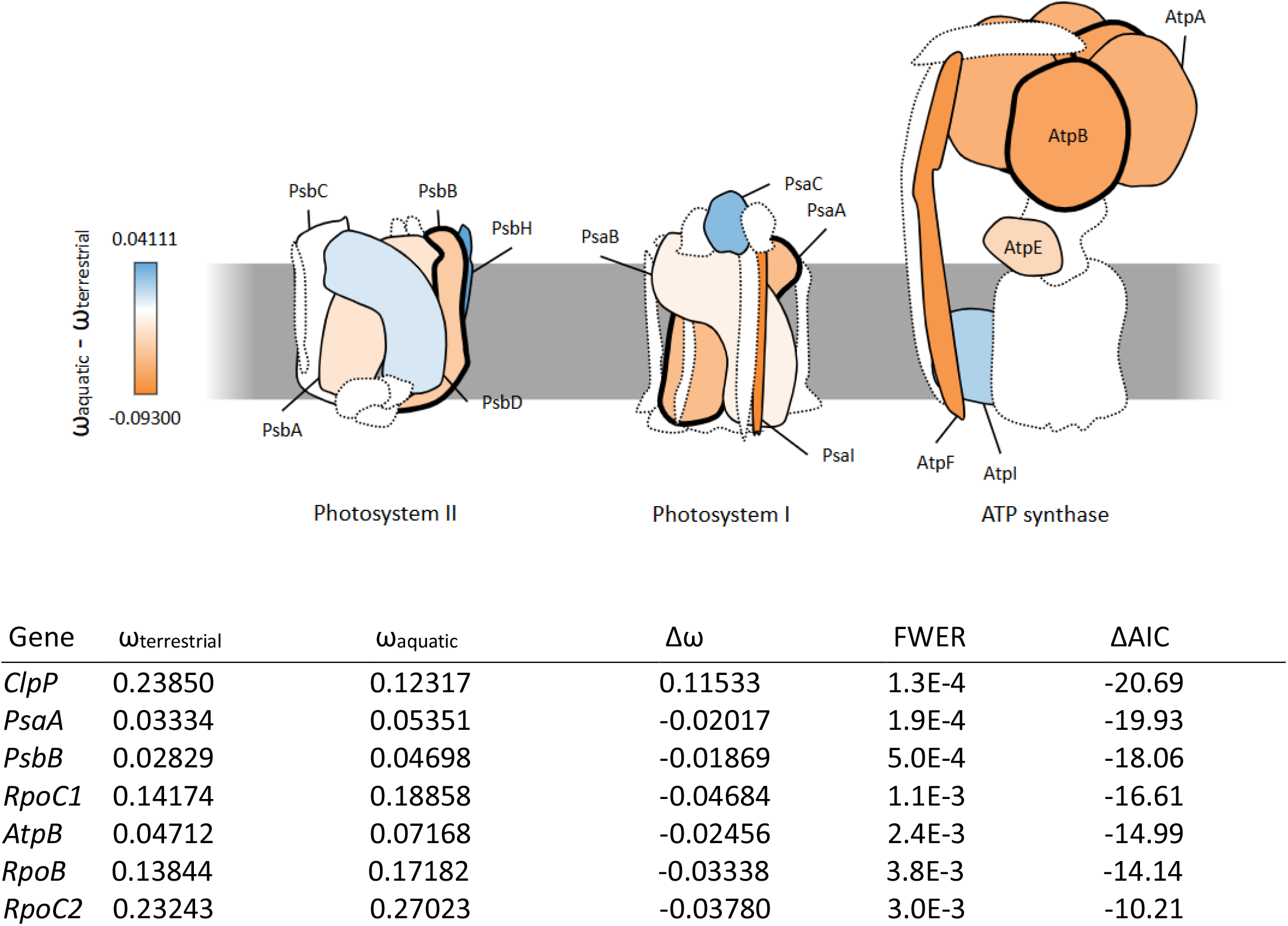
Evolution of ω in chloroplast photosystems and ATP synthase between aquatic and terrestrial plants. Blue indicates lower ω in aquatic plants, and orange indicates lower ω in terrestrial plants. Dotted lines indicate that ω could not be calculated, and thick lines indicate significant differences between aquatic and terrestrial ω, *i*.*e*. differences associated with a Family-Wise Error Rate (FWER) < 0.05 and a difference in Akaike Information Criterion (ΔAIC) < −2. The bottom table shows all significant results.

Of these seven genes, two (*PsaA* and *PsbB*) are major components of photosystems (PS) and three (*RpoC1, RpoC2* and *RpoB*) are DNA-directed RNA polymerase subunit. In addition to photosystems, the other two genes found are also involved in photophosphorylation, with *AtpB* being a subunit of ATP synthase and *ClpP* being involved in the degradation of cytochrome b_6_f [19]. In addition to these analyses, we have performed a comparison of orthogroup expansions and contractions between aquatic and terrestrial plants on the same sets of genomes. The results obtained in this manner are not very informative and more subject to interpretation. Thus, they are not discussed here but are nevertheless presented in Figure S5, with additional information in Supplementary methods.

## Discussion

The lifestyle of terrestrial plants is subject to numerous constraints that have shaped their physiology and morphology from their terrestrialization to the present day. Here, this is evidenced by the genes associated with stronger selection pressure in terrestrial plants, mainly the *DTX* genes. In *A. thaliana*, DTX play a role in the export of toxic compounds and in apical dominance and leaf initiation [25,26]. Stronger selection pressure was also detected in terrestrial plants for COBRA-like genes, which are involved in cellulose structure and organization [27,28]. A change in cellulose organization is thought to have played a key role in terrestrialization, as it may have favored directed growth or increased rigidity [29]. Interestingly, most aquatic plants do not organize themselves around a main axis but rather rely on basal growth, which may explain a relaxation of selection pressure on this mechanism, while it provides a competitive advantage in access to light for terrestrial species. In contrast, few genes related to plant structure or development are under selective pressure in aquatic plants. Stronger selection pressure on pectin-catabolism-related *PME* genes and enrichment in GO terms “Programmed Cell Death” and “Aerenchyma formation” might together highlight formation of aerenchyma [30–32]. Other physiological adaptations found in aquatic plants is related to genes encoding blue-light-receptor cryptochromes and genes more or less directly linked to them. Since blue light penetrates the deepest into the water [33], it can be assumed that an adaptation of cryptochromes in aquatic plants allows them to better perceive blue light for various metabolic processes. Many genes related to oxidative stress and ROS homeostasis were also observed among the genes selected by sPLS-DA. In particular, orthologs of the *A. thaliana MIOX* and *GULLO* genes, involved in ascorbate synthesis [11,12], are associated with stronger selection pressure in aquatic plants, as are orthologs of *APX* genes, which encode ascorbate peroxidases that use ascorbate to scavenge hydrogen peroxide [34]. As ascorbate is an extremely important molecule for ROS homeostasis, our results show that adaptation to an aquatic lifestyle is primarily an adaptation to regular oxidative stress.

It is assumed that terrestrial environments are generally more diverse than aquatic environments [35,36]. For these reasons, terrestrial plants must adapt to efficiently uptake nutrients, which is marked by an enrichment in major genes related to sulfate, phosphate assimilation and to autophagy-under-starvation-related genes. Recent studies have highlighted the importance of mycorrhizal symbiosis in terrestrialization [37], and thus also the increased importance of nutrient assimilation for the terrestrial lifestyle. In comparison, aquatic plants can draw nutrients from the water column, which can be renewed and homogenized by water flows to benefit plant growth [38,39]. Some aquatic plants also appear to use other nitrogen sources than nitrate [40].

For aquatic plants, potassium transporters were associated with high selection pressure, but they might be linked to potassium loss in hypoxic environments, which can normally lead to cell death [41], rather than starvation. In addition to nutrient assimilation, many external stimuli must be perceived in a terrestrial heterogeneous environment, which is evident in our results with the enrichment of terms related to signal transduction, endosomes, intracellular transport, and cellular communication. These results suggest that terrestrial plants are more receptive to their environment and faster to adapt to it. Concomitantly, an expansion of signaling-related genes is thought to have participated in the pre-adaptation of algae to terrestrialization [42,43].

The results of ω analyses are difficult to interpret because a high ω is supposed to indicate relaxed or positive selection pressure. Here, the aquatic plants descended from terrestrial ancestors, and form a paraphyletic group. For these reasons, a slightly higher ω_aquatic_ than ω_terrestrial_ could in fact show convergent adaptations to the aquatic lifestyle rather than a relaxation of selection pressure. Our most informative and significant results were obtained from chloroplastic genes: *PsaA* and *PsbB* genes, two major components of photosystems, evolve differently in terrestrial and aquatic plants. Here, we propose that the evolution of these genes optimizes the harvesting of certain wavelengths depending on the environment, a known adaptation in PsbB [44]. Structural changes in photosystems could also reduce redox potentials to avoid electron leakage and subsequent oxidative stress in aquatic plants, or increase it to accelerate photophosphorylation and take advantage of higher CO_2_ availability in terrestrial plants. Finally, the strongest differences between ω are mainly found within the ATP synthase complex, which may highlight adaptations to proton gradient magnitude.

Only one gene, *ClpP*, was associated with lower ω in aquatic plants, indicating a likely aquatic lifestyle adaptation. ClpP is notably involved in the degradation of defective cytochrome b_6_f. Cytochrome b_6_f has many roles within the photosynthetic chain, that have been summarized by Malone *et al*. [45]. It is notably involved in regulating the photosynthesis according to the linear electron transfer rate and in preventing short-circuits of the mitochondrial Q-cycle, as cytochrome b_6_f is also a component of mitochondria, which produces ROS under hypoxia [46]. Thus, renewal of defective cytochromes b_6_f may be a key mechanism for aquatic plants to limit ROS production during O_2_ level fluctuations.

Our results show that the terrestrial lifestyle appears to be more restrictive overall than the aquatic lifestyle, to the point that one might wonder why plants bothered to leave the water in the first place. One hypothesis is that pre-adapted algae occupied the vacant ecological niche of dry land opportunistically. However, we emphasize that the terrestrial lifestyle appears to be so constraining that factors other than opportunism may have played a role. Our results show that the main constraint of the aquatic lifestyle appears to be oxidative stress, and it is possible that access to an environment with stable and high amounts of O_2_ and CO_2_ favored the emergence of terrestrial plants. Indeed, recent estimates date plant terrestrialization to ~497 million years ago [47], a supposed period of a major ocean anoxia event associated with pulses of increased atmospheric oxygen and high CO_2_ levels [48,49]. This allowed to accelerate the energy mechanism, notably through adaptations of the chloroplast electron transport chain. Oceanic anoxia may also have played a role, although it is not certain that this event affected freshwater bodies from which embryophytes are thought to have arisen [50]. Alternatively, a fossil of Charophyceae living in an Ordovician marine environment was discovered in 2025 [51], and it has been proposed that adaptation to fresh water occurred after terrestrialization [52]. In both cases, it is possible that terrestrialization of plants was promoted by an anoxia event, in addition to the opportunistic occupation of a vacant ecological niche.

By comparing a set of aquatic and terrestrial plants, we highlighted the different main mechanisms of return to water and also identified certain mechanisms essential to terrestrial plants. We also proposed a hypothesis explaining the terrestrialization of plants despite the apparent complexity of adapting to such a lifestyle. Due to limited data, our study focuses primarily on angiosperms, and some of these mechanisms could be more or less angiosperm-specific, although the results obtained with *M. polymorpha* do not appear to differ fundamentally from those of other terrestrial plants. Hopefully, greater availability of non-angiosperm embryophyte genomes in the near future could help to elucidate certain stress-related mechanisms that were not pointed by our analyses. Data on intra-species genetic variability in non-angiosperms are even scarcer, but could also reveal selective pressures that we have not yet been able to identify.

## Methods

### Material

The complete list of the 56 sequences and genomes used for population genetics calculations is available in Table S5, with data names, species names, authors, and references, as are the eight genetic variability files. The complete list of the 129 sequences and genomes used for ω calculations is available in Table S6 with the same information, as well as the genome type (chloroplast or nuclear). Of all sequences used, only *R. penicilliatus* genome was not already publicly available. *R. penicilliatus* specimen was collected in Hiis, Occitanie, France (43.135919, 0.103214) by François Prud’Homme from the Conservatoire Botanique National Pyrénées et Midi-Pyrénées in accordance with all relevant institutional, national, and international guidelines and legislation. It was sequenced at the Centre National de Ressources Génomiques Végétales (CNRGV, Castanet-Tolosan, France). *R. penicilliatus* genome is currently being processed by NCBI (PRJNA1358412) and the complete set of genetic variation data generated for this study, are available in the Data From Comparative Genomic Insights into the Evolution of Aquatic and Terrestrial Adaptations in Plants Figshare repository (https://doi.org/10.6084/m9.figshare.30998356, 10.6084/m9.figshare.30998356).

### Genetic population statistics

Genetic variation data for *N. peltata, P. oceanica, R. bungei* and *P. crassipes* were generated by mapping various sequencing data to different genomes (Table S5). Furthermore, inference of ancestral alleles was done for *A. thaliana, B. distachyon* and *M. truncatula* to compute Zeng’s E and Way and Fu’s H (see Supplementary methods for more information). Note that *R. bungei* was mapped on a the newly-sequenced *R. penicilliatus* genome. Tajima’s D, Zeng’s E and Way and Fu’s H were calculated as described in [20], but considering only the coding parts of the genes. The significance of differences in mean statistics between terrestrial and aquatic plants was then measured with Wilcoxon-Mann-Whitney tests. The NS/S ratio was calculated by dividing the number of non-synonymous polymorphic sites with the number of synonymous polymorphic sites, determined using SNPEff. When no structural annotation was available (*P. oceanica, S. stoloniferum, P. crassipes, R. penicilitaus*), it was generated by masking the genomes with Red and predicting genes and CDS positions with BRAKER (for version and references of all the bioinformatics tools used, see Table S7). For this step, the OrthoDB v11 *Viridiplantae* set [21] was used as input protein sequences. Orthogroups were determined with sets of protein encoded by the longest transcript for each gene in each species and then using OrthoFinder with a fixed species tree and the -M msa option. The fixed tree was generated by an initial run of OrthoFinder, followed by a manual check between the structure of the OrthoFinder-generated tree and the known phylogeny of embryophytes. When paralogs were present within an orthogroup, the values of each of their statistics were averaged. To avoid biases related to large variations between paralog number, the number of paralogs for each species inside each orthogroups was computed. Orthogroups with more than one outlier identified by the interquartile range (IQR) method, which detects outliers as values falling below Q1 − 1.5 × IQR or above Q3 + 1.5 × IQR, or with a coefficient of variation > 1, were thus removed. Each statistics was scaled within each species and an sPLS-DA was then performed for each statistics (Tajima’s D and NS/S) with the R mixOmics package, keeping only the 100 more discriminant variables (with parameter keepX = c(100,100)). Finally, enrichments were made with the STRINGdb package from the *A. thaliana* genes contained in the orthogroups selected by the sPLS-DAs. Dispensability of the enriched GO terms was calculated with the REVIGO web application.

### ω comparisons

New sets of orthogroups were inferred from numerous genomes (Table S6). For each orthogroup, we removed parts of OrthoFinder’s gene trees corresponding to unresolved relationship using the Python library ETE 3. Genes that were removed from the trees were then removed from the orthogroups, and we kept only the orthogroups where at least seven aquatic and seven terrestrial species were still present to assure a good representativeness of both types of plants. The protein sequences of each orthogroup were aligned using MAFFT. Poorly aligned sequences were filtered with trimAl with option -seqoverlap 80 and alignments were then transformed into codon-based alignments with the PAL2NAL program with the -nogap option. The resulting alignments were filtered to keep only those larger than 90 bases. Then, CODEML was used to calculate, on each orthogroup, the global ω (model M0) and a different ω for aquatic plants and terrestrial plants (model M2). To select the most significant results, only the orthogroups with an Akaike Information Criterion (AIC) [22] difference lower than −2 between the M2 and M0 models were included. The significance of the results was measured by Likelyhood Ratio Test (LRT) [23], and the resulting p-values were transformed into Family-Wise Error Rate (FWER) using Holm’s method [24], as it is more stringent than False Discovery Rate (FDR) methods. Finally, only orthogroups with a FWER < 0.05 were selected.

## Acknowledgments

We are thankful to Mr. Hung Manh Nguyen and Mr. Masayuki Maki for providing data for *P. oceania* and *N. peltata*, respectively, as well as to Mr. Evan Ernst for allowing us to use data from lemna.org, even if these were not ultimately used in the final version of this article. We also thank Maxime Bonhomme and Jean Keller from the Laboratoire de Recherche en Sciences Végétales for their advice on calculating Tajima’s D, Zeng’s E, Way & Fu’s H, and gene expansions and contractions. We’re also grateful to Francois Prud’homme from the Conservatoires Botaniques Nationaux Pyrénées et Midi-Pyrénées for his help to collect *R. penicilatus* material, and to Caroline Callot and William Marande from the Centre National de Ressources Génomiques Végétales for genome sequencing and assembling.

## Data availability statement

*R. penicilliatus* genome is currently being processed by NCBI (PRJNA1358412) and the complete set of genetic variation data generated for this study, are available in the Data From Comparative Genomic Insights into the Evolution of Aquatic and Terrestrial Adaptations in Plants Figshare repository (https://doi.org/10.6084/m9.figshare.30998356, 10.6084/m9.figshare.30998356). All other data used come from public databases (see Table S5 and Table S6).

## Author contributions

S. Cabanac did all the investigation and analysis process, and prepared figures. All authors contributed to the conceptualization and wrote the main manuscript text

---

**Figure S1.**
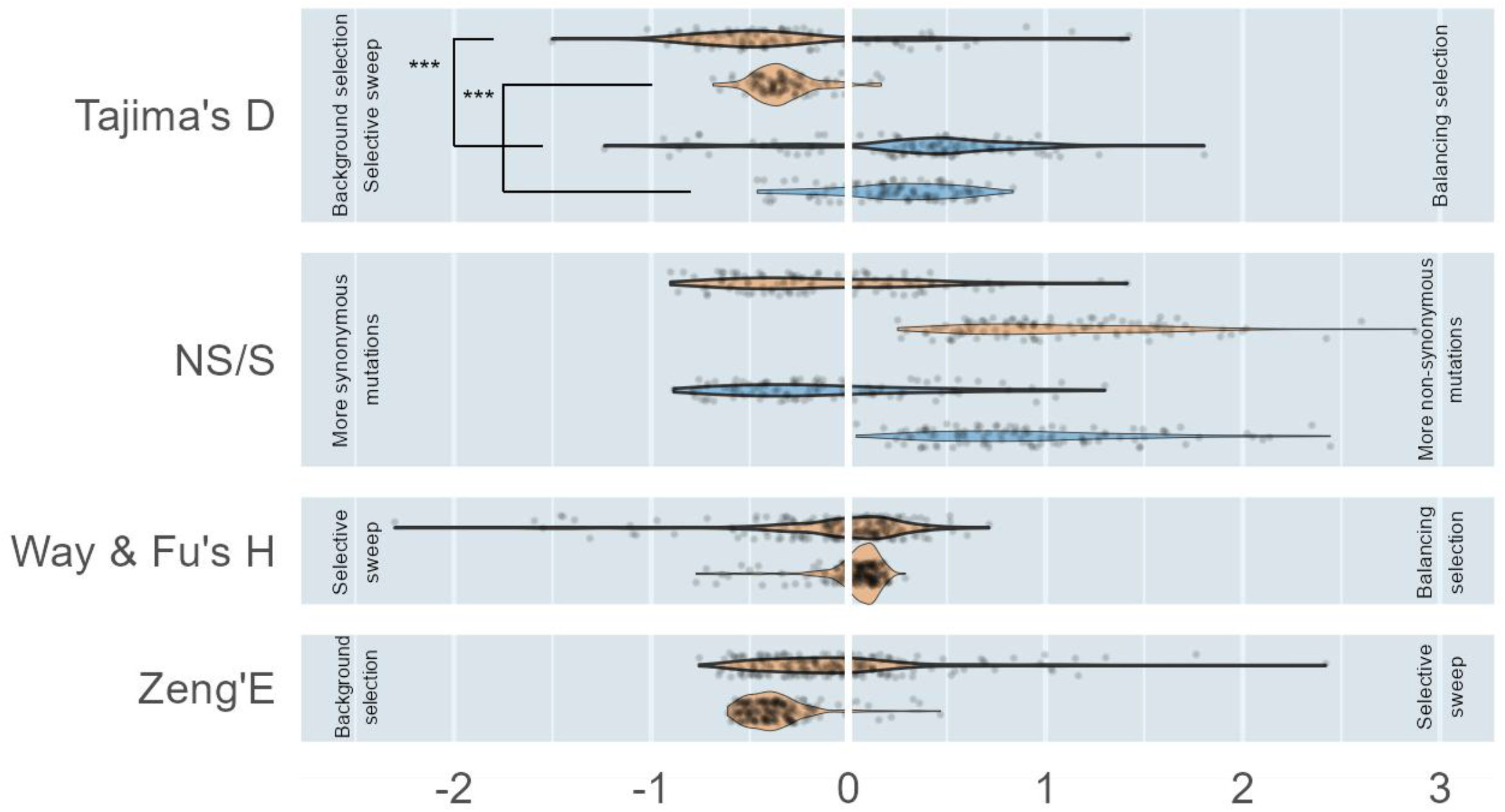
Distribution of values obtained for different population genetic statistics. Values are the mean values for terrestrial (orange) and aquatic (blue) plants within the orthogroups selected by sPLSDA_D_ and sPLSDHA_NS/S_. The violins surrounded by a thick line were obtained with the scaled values within each species. *** indicate a significant difference between the means (P-value < 0.001). The values for Zeng’s E and Way & Fu’s H are those associated with genes obtained by the sPLS-DA using either Tajima’s D or NS/S values, indifferently

**Figure S2.**
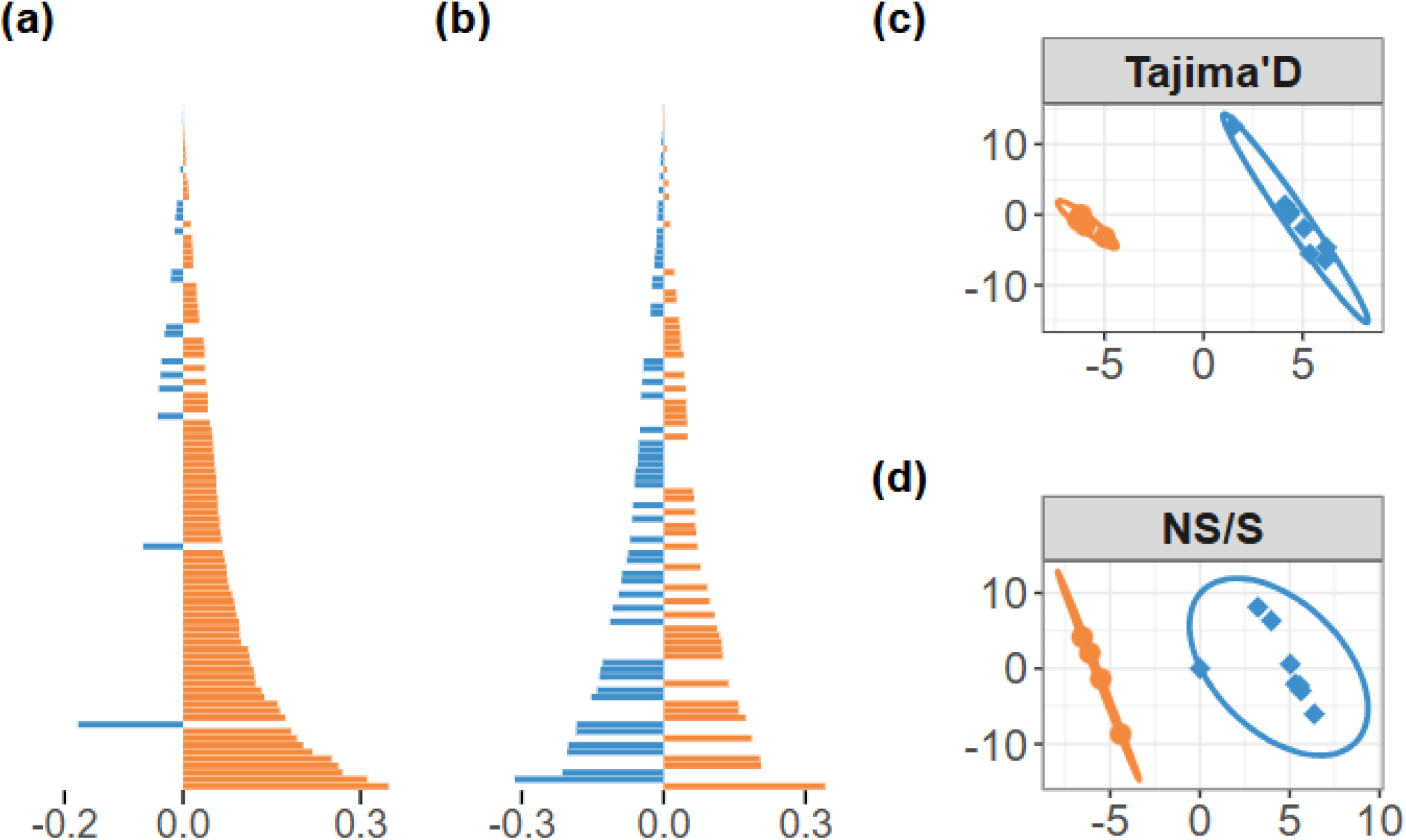
sPLS-DA results obtained from scaled NS/S and Tajima’s D values, between a set of terrestrial and aquatic plants. (a) Overall trend of the results obtained from Tajima’s D values, representing the loadings on the X-variate-1 of the 100 orthogroups associated with the strongest loadings. Blue bars indicate orthogroups where aquatic plants are associated with lower average scaled Tajima’s D values, and orange bars indicate those where terrestrial plants are associated with lower average scaled Tajima’s D values. (b) Same as (a) with NS/S values. (c) Representation of terrestrial (orange) and aquatic (blue) species in the space defined by the first two axes from the sPLS-DA obtained from Tajima’s D or (d) NS/S values.

**Figure S3.**
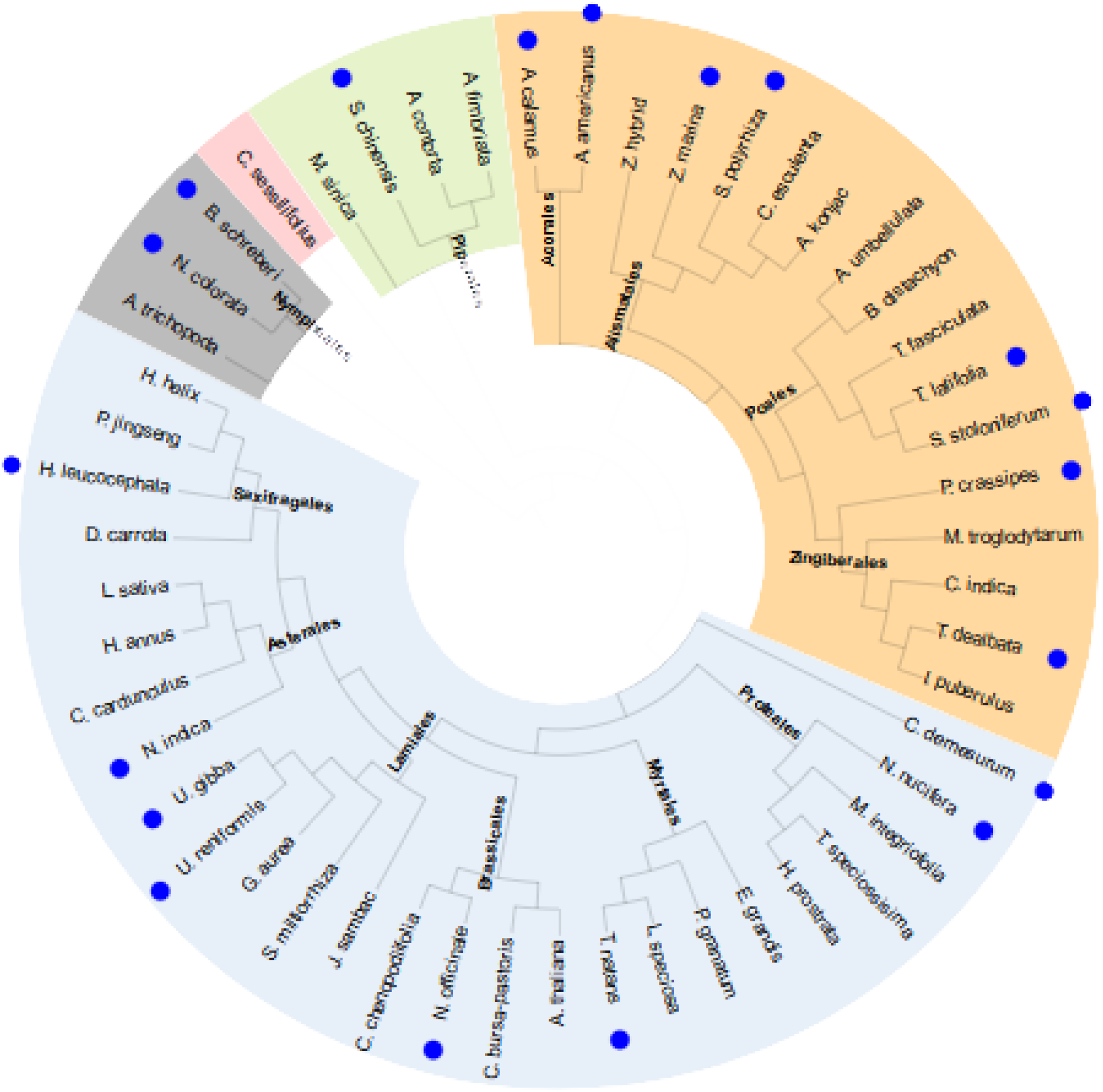
Phylogenetic tree of species used for gene expansion comparison, according to NCBI and APG IV taxonomies. Aquatic species are marked with a blue dot. Orders are written on the branches, and colored backgrounds indicate major clades. Gray = basal angiosperms, pink = Chloranthales, green = Magnoliids, orange = Monocots, blue = Eudicots.

**Figure S4.**
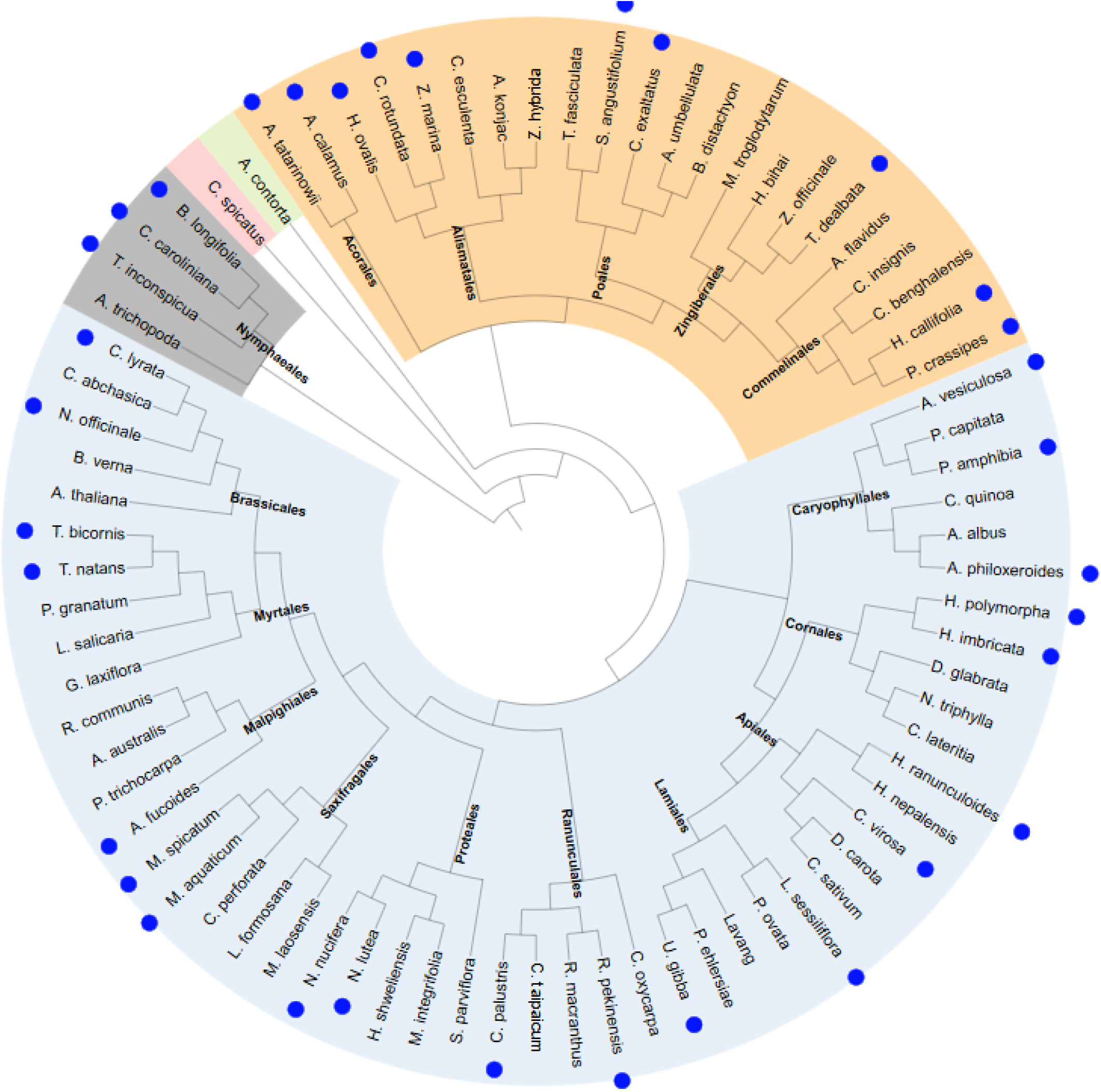
Phylogenetic tree of species used for chloroplast gene evolution comparison, according to NCBI and APG IV taxonomies. Aquatic species are marked with a blue dot. Orders are written on the branches, and colored backgrounds indicate major clades. Gray = basal angiosperms, pink = Chloranthales, green = Magnoliids, orange = Monocots, blue = Eudicots.

**Figure S5.**
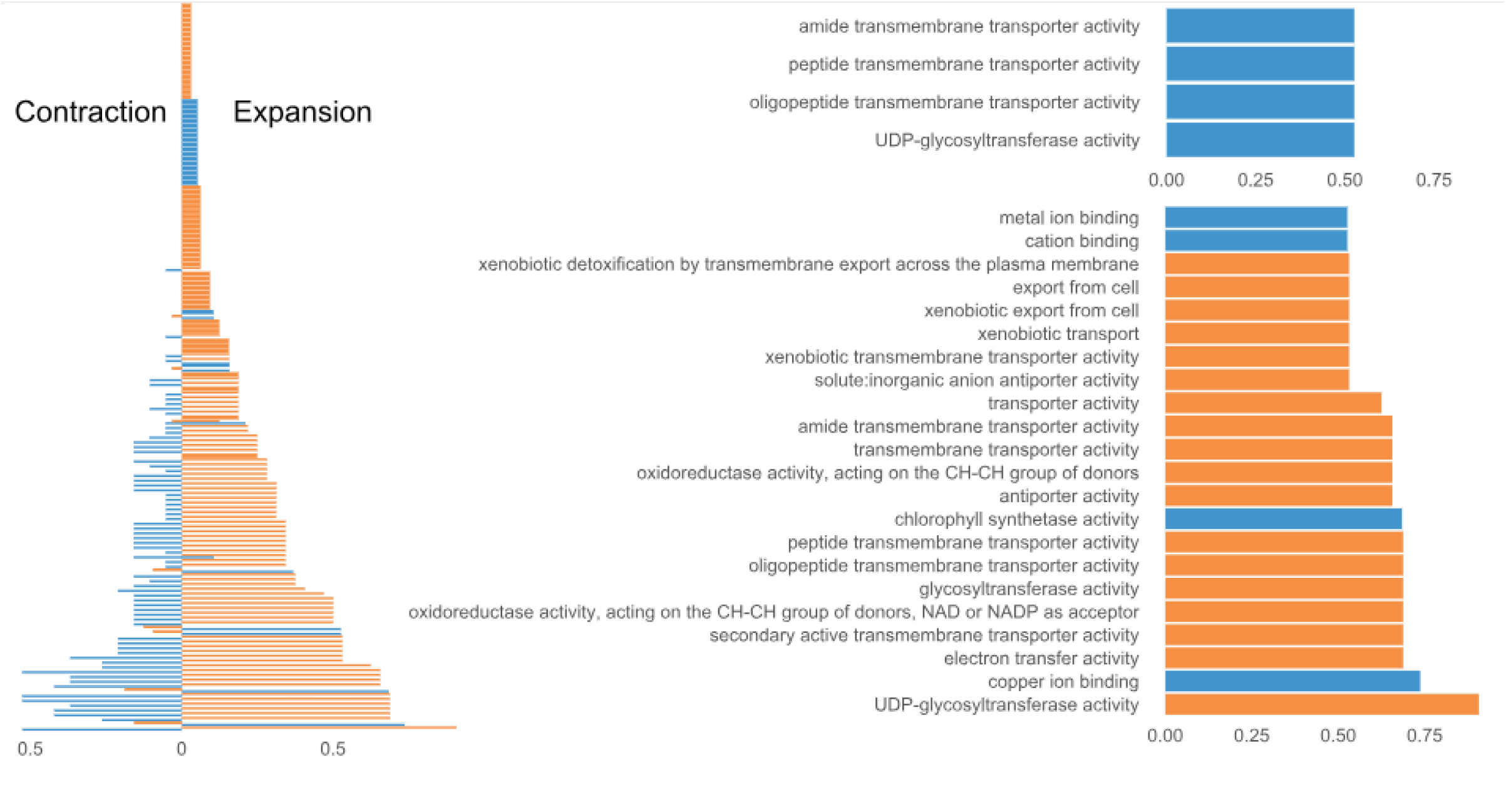
Frequency of GO term enrichments among genes selected by sPLS-DA from the number of orthologs inside each orthogorup. (a) Global contraction and expansion trends among aquatic plants (blue) and terrestrial plants (orange). The X-axis shows the ratio of species for which the term is enriched. (b) Zoom in on the most contracted GO terms. (c) Zoom in on the most expanded GO terms. Note that for clarity, only GO terms with a frequency greater than 0.5 are shown.

